# Pollutants in Hong Kong Soils: Organochlorine Pesticides and Polychlorinated Biphenyls

**DOI:** 10.1101/2020.02.16.951541

**Authors:** M.K. Chung, R. Hu, K.C. Cheung, M.H. Wong

## Abstract

Organochlorine pesticides (OCPs) and polychlorinated biphenyls (PCBs) were investigated in 138 soil samples collected in Hong Kong. Results showed that within the selected OCPs, only DDT and its metabolites (DDD and DDE) were frequently detected. Of 138 soil samples, 18% were non-detected for any DDT residues, while 25% were contaminated sporadically with DDT only (non-detected to 1090 µg kg^−1^) and 57% contained various combinations of DDT, DDD and DDE (2.03 to 1118 µg kg^−1^). In contrast, total PCBs (∑66 congeners) contamination was distributed more evenly (0.22 to 154 µg kg^−1^) than DDTs, but serious contamination was found in industrial areas and soils collected near highways. Concentrations of 7 indicator PCBs ranged between non-detected to 34.5 µg kg^−1^. The ratios of DDT/metabolites were typically greater than 1, thus suggesting recent application of DDT. Medium-range deposition from industrial areas within as well as away from the city is also suspected to be the origins of DDTs and PCBs found in Hong Kong soils. The concentrations of DDTs and PCBs in all soil samples did not exceed the recommended values in soil quality guidelines except 3 samples for DDT from locations far away from densely populated districts such as Tuen Mun and Tin Shui Wai. Therefore, DDTs and PCBs were not of significant concern in terms of their impacts on public health and environment.

## 1. Introduction

Polychlorinated biphenyls (PCBs) and a number of organochlroine pesticides (OCPs) such as dichlorodiphenyltrichloroethane (*p,p’*-DDT) and hexachlorocyclohexanes (HCHs) are candidates of designated 12 Stockholm Convention persistent organic pollutants (POPs). These anthropogenic chemicals, unlike other nature contaminants that can be degraded by physiochemical or biological means, are much more persistent in the environment. Their potential carcinogenicities and the possibility of resulting more toxic metabolites have spotlight attention of the general public in recent years (Connell *et al.*, 2003; Megharaj *et al.*, 1999).

PCBs were first commercially produced in 1929 (Mai *et al.*, 2005) and cumulative production has been estimated as much as 1.2 million tonnes around the world (Harrad *et al.*, 1994). The primary uses of PCBs included congener-mixture in dielectric fluids, flame retardants, and industrial lubricant fluids in transformers and capacitors (Schuhmacher *et al.*, 2004). They are organochlorine chemicals that have 209 congeners. Among these congeners, 12 are especially dangerous for their dioxin-like (DL) activities (van den Berg *et al.*, 1998). Their residues have been found in the environment and have been either banned or restricted on production or usage in many countries since 1970s (Harrad *et al.*, 1994).

DDT was first synthesized in 1874 and the discovery of DDT’s insecticidal activity by Paul Müller in 1939 subsequently led to his award of the Nobel Prize (Carson *et al.*, 1962). After World War II, it was applied on agricultural crops world wide and stimulated the synthesis and development of other organochlorine pesticides. However, later evidences showed its detrimental effects on non-target organisms (Carson *et al.*, 1962; Christen, 1999), leading to the restriction on its production and usage in many countries. Nevertheless, several tropical and subtropical countries are being exempted for using DDT for public health purpose to control the spread of malaria (World Wildlife Fund, 2004).

In Asia, Japan and Korea have banned the use of DDT in the 1970s (Phillips and Tanabe, 1989), whereas DDT production was prohibited in China in 1983 (Wolfe *et al.*, 1984). PCBs have been banned or regulated in China in early 1980s (Mai *et al.*, 2005). Nevertheless being officially regulated for more than 20 years, PCBs, DDTs and other OCPs such as HCH were still being detected in various environmental compartments throughout China (Ding *et al.*, 2005; Fung *et al.*, 2004; Mai *et al.*, 2005; Wu *et al.*, 1999; Yuan *et al.*, 2001), but overall the PCB levels were relatively low and serious contaminations were highly restricted in area such as storage locations of PCBs (Xing *et al.*, 2005). In Hong Kong, the production and use of DDT have been banned since 1988, however, there is no legislation to ban the production of PCBs as no record of such industrial activity by the Hong Kong government. Instead, historical equipments with PCBs such as transformers and capacitors are being phased out voluntarily by their owners (Hong Kong Environmental Protection Department, 2002).

In the Pear River Delta (PRD) where Hong Kong is located, Lingding Yang was reported to be one of the hotspot sites polluted with OCPs in the Pearl River Estuarine (Fang, 2004), which might transport pollutants to coastal areas of Hong Kong. Contamination of PCBs and DDTs in mussels farmed in Hong Kong was reported as early as 1990s (Phillips, 1989). DDT levels in human milk collected from both Guangzhou and Hong Kong were generally higher when compared with 19 countries (Wong *et al.*, 2005). Recent studies indicated that several freshwater fishes cultivated in fish ponds around the PRD and purchased from local markets all contained DDTs (Cheung *et al.*, 2006), and in particular the Mandarin fish (*Siniperca kneri*) are of actual concern, which could contain DDTs up to 4.3 times higher (62 µg kg^−1^ wet wt.) than the recommended value set by USEPA (14.4 µg kg^−1^ DDTs, wet wt.) (Kong *et al.*, 2005).

The major objectives of this paper are to study the contemporary levels of edaphic OCPs and PCBs and assess their potential risks to general public in Hong Kong. It is hoped that the data would serve as a valuable reference for redeveloping of some areas which were affected by various industries in the past, as well as fulfilling part of the obligation imposed by the Stockholm Convention on POPs and the duty of monitoring POPs in the environment. In addition to these, potential sources of OCPs and PCBs were also discussed.

## 2. Materials and Methods

### 2.1 Sampling, Preparation and Analysis

Ten land use categories had been designated to reveal the pollution impacts from various human activities. These included urban park, greening area, country park, rural area, restored landfill, agricultural farmland, orchard farm, crematorium, industrial area and nearby highway. There were totally 138 composite soil samples collected in early 2003 at various locations in Hong Kong. Surface soils (0-5 cm depth) were taken by using a stainless steel soil core with the uppest organic vegetative materials were removed in advance. Samples were air-dried and sieved through a 2-mm mesh.

Extraction of OCPs and PCBs from soil samples were performed according to the US EPA Standard Method 3540C (U.S. Environmental Protection Agency, 1996a). Briefly, 10 g of soil sample was transferred into a soxhlet apparatus and extracted by 80 ml acetone (pesticide grade, Tedia) and dichloromethane (DCM) (pesticide grade, Tedia) mixture (1:1, v:v) for 18 h. Florisil columns were used to minimize interferences to target compounds.

Fifteen OCPs were analyzed: α-HCH, β-HCH, δ-HCH, heptachlor, aldrin, heptachlor expoxide, endosulfan I, endosulfan II, dieldrin, *p,p’*-DDE, endrin, *p,p’*-DDD, endrin aldehyde, endosulfan sulphate and *p,p’*-DDT. For PCBs, 66 of the congeners were quantified. Using the IUPAC nomenclature, they are 1, 2, 3 (total mono-PCBs), 4, 6, 8, 9, 15 (total di-PCBs), 16, 18, 19, 20, 22, 25, 27, 28, 29, 34 (total tri-PCBs), 40, 42, 44, 47, 52, 56, 66, 67, 69, 71, 74 (total tetra-PCBs), 82, 87, 92, 93, 99, 101, 105, 110, 118, 119 (total penta-PCBs), 128, 134, 136, 138, 144, 146, 147, 151, 153, 157, 158 (total hexa-PCBs), 173, 174, 177, 179, 180, 187, 190, 191 (total hepta-PCBs), 194, 195, 199, 203 (total octa-PCBs), 206, 207, 208 (total nona-PCBs) and 209 (total deca-PCBs). ∑PCBs is defined as the sum of the concentration of 66 congeners. Standards for OCPs and PCBs congeners mixtures were purchased from ChemService Inc. and AccuStand Inc. respectively.

The analytical methods were based on using GC-MS (Gas Chromatrography-Mass Spectrometry) instrument (U.S. Environmental Protection Agency, 1996b). In short, OCPs and PCBs were quantitatively analyzed by a Hewlett Packard (HP) 6890 GC system equipped with an HP 5973 mass selective detector (MS) and a 30m × 0.25mm × 0.25µm DB-5 capillary column (J & W Scientific) with helium as the carrier.

Limit of detection (LOD) for OCPs was: 2 µg kg^−1^ for *p,p’*-DDD and *p,p’*-DDE, 20 µg kg^−1^ for DDT and 10 µg kg^−1^ for the rest of OCPs. LOD for PCBs varied, and a universal safe value for mono to penta PCBs was set to 0.5 µg kg^−1^ while 1 µg kg^−1^ for the rest of the PCBs.

### 2.2 Quality Control

Laboratory analytical blank and Certified Reference Material (CRM) CRM105-100 (Resource Technology) and HS-2 (Institute for Marine Biosciences) were included in every 2 batches (14 samples) of soxhlet extraction to assess the recoveries and performance for measurement of OCPs and PCBs respectively. None of the analytical blanks were found to have detectable contamination of the monitoring OCPs and PCBs. Average individual OCPs recoveries were: 88 ± 3% (*p,p’*-DDT), 92 ± 6% (*p,p’*-DDD), 86 ± 2% (*p,p’*-DDE), 67 ± 4% (dieldrin), 64 ± 7% (endosulfan I), 72 ± 6% (endosulfan II), 59 ± 9% (endrin). While average individual PCBs recoveries were: 77 ± 3% (IUPAC 101), 89 ± 5% (IUPAC 138), 87 ± 5% (IUPAC 151), 103 ± 9% (IUPAC 153), 76 ± 12% (IUPAC 180), 82 ± 3% (IUPAC 194), 86 ± 10% (IUPAC 199), 78 ± 9% (IUPAC 209). Mean recovery for OCPs and PCBs were > 75% and >84% respectively and all samples were not corrected.

### 2.3 Data Analyses

Statistical analyses were calculated using Statistica (version 6.0, StaSoft). Kriging maps were developed using SADA (version 4.0, University of Tennessee) and graphical plots of data were produced by either Statistica or Sigmaplot (version 8.0, Systat). Not detected values were substituted with half of LOD only for descriptive statistics.

## 3. Results and Discussion

### 3.1 Spatial values

In general, most OCPs were not detected except DDTs, and their mean concentrations are listed in Table 1. Two locations were found to have endosulfan sulfate at concentration about 12 µg kg^−1^ while heptachlor epoxide was found in 1 soil sample. This finding is in HK is in line with the observation that DDTs were the mostly predomonant OCPs in tropical Asia (Kannan *et al.*, 1995). Uneven distribution of DDTs was observed, with 18% of soil samples showing no detectable DDTs. Areas near highways had higher concentrations of DDT when compared to other land use categories (median 77.8 µg kg^−1^). DDT concentrations decreased according to the following pattern: areas near highways > urban park > crematorium > orchard farm > rural area > agricultural farmland > greening area > industrial area > country parks > restored landfills. The variation in concentrations was greatest for DDT, ranged from not detected to 1000 µg kg^−1^, whereas not detected to around 200 µg kg^−1^ for DDD and DDE.

**Table 1.** Mean concentrations of DDTs and PCBs of soils in Hong Kong. Units are in µg kg^−1^ except for the WHO-TEQ PCBs.

| Classified soil categories | Sample no. | DDT |  |  | DDD |  |  | DDE |  |  | Total DDTs |  |  |  |  |  |
| --- | --- | --- | --- | --- | --- | --- | --- | --- | --- | --- | --- | --- | --- | --- | --- | --- |
|  |  | M* | Medi* | Range | M* | Medi* | Range | M* | Medi* | Range | M* | Medi* | Range |  |  |  |
|  | Concentration in soils (µg/kg) |  |  |  |  |  |  |  |  |  |  |  |  |  |  |  |
| Urban park | 39 | 39.7 | 34.2 | [N.D. - 160] | 4.4 | 4.4 | [N.D. - 27.9] | 2.5 | 2.3 | [N.D. - 8.3] | 2.5 | 2.3 | [N.D. - 165] |  |  |  |
| Greening area | 14 | 24.4 | 14.3 | [N.D. - 162] | 3.8 | 4.5 | [N.D. - 6.4] | 1.6 | 2.1 | [N.D. - 4.1] | 1.6 | 2.1 | [N.D. - 169] |  |  |  |
| Country park | 9 | 18.1 | 2.0 | [N.D. - 51] | 0.8 | 0.2 | [N.D. - 3.5] | 0.2 | 0.2 | [N.D. - 1.7] | 0.2 | 0.2 | [N.D. – 51] |  |  |  |
| Rural area | 19 | 35.9 | 19.4 | [N.D. - 252] | 1.3 | 0.2 | [N.D. - 4.4] | 0.7 | 0.2 | [N.D. - 2.4] | 0.7 | 0.2 | [N.D. - 253] |  |  |  |
| Restored landfill | 11 | 4.0 | 2.0 | [N.D. - 31.5] | 2.5 | 0.2 | [N.D. - 7.7] | 1.3 | 0.2 | [N.D. - 3.8] | 1.3 | 0.2 | [N.D. – 43] |  |  |  |
| Agricultural farmland | 9 | 42.0 | 19.0 | [7.8 - 125] | 3.5 | 3.6 | [N.D. - 7.5] | 3.8 | 2.1 | [0.75 - 15.7] | 3.8 | 2.1 | [8.71 - 149] |  |  |  |
| Orchard farm | 5 | 35.0 | 19.8 | [16.3 - 67.2] | 6.3 | 3.9 | [N.D. - 19.4] | 21.9 | 1.9 | [N.D. - 103] | 21.9 | 1.9 | [20.2 - 190] |  |  |  |
| Crematorium | 10 | 22.0 | 21.1 | [N.D. - 54.1] | 1.9 | 2.4 | [N.D. - 3.6] | 1.0 | 1.2 | [N.D. - 2.1] | 1.0 | 1.2 | [4.03 – 55] |  |  |  |
| Industrial area | 18 | 257.0 | 12.7 | [N.D. - 1090] | 25.5 | 2.6 | [N.D. - 215] | 12.8 | 0.2 | [N.D. - 108] | 12.8 | 0.2 | [N.D. - 1118] |  |  |  |
| Nearby Highway | 4 | 103.0 | 77.8 | [22.6 - 232] | 1.4 | 0.2 | [N.D. - 5.4] | 0.8 | 0.3 | [N.D. - 2.6] | 0.8 | 0.3 | [23 - 241] |  |  |  |
| Classified soil categories | Sample no. | PCB 105 (DL) |  |  | PCB 118 (DL) |  |  | PCB-157 (DL) |  |  | Total 3 DL PCBs |  |  | ng PCB WHO-TEQ kg-1 |  |  |
|  |  | M* | Medi* | Range | M* | Medi* | Range | M* | Medi* | Range | M* | Medi* | Range | M* | Medi* | Range |
|  | Concentration in soils (µg/kg) |  |  |  |  |  |  |  |  |  |  |  |  |  | Concentration (ng/kg) |  |
| Urban park | 39 | 0.11 | N.D. | [N.D. - 0.68] | 0.18 | 0.1 | [N.D. - 1.05] | 0.05 | N.D. | [N.D. - 0.4] | 0.34 | 0.1 | [N.D. - 2] | 0.05 | 0.02 | [N.D. - 0.36] |
| Greening area | 14 | 0.15 | 0.1 | [N.D. - 0.8] | 0.2 | 0.1 | [N.D. - 1.24] | 0.04 | N.D. | [N.D. - 0.16] | 0.39 | 0.2 | [N.D. - 2.12] | 0.05 | 0.03 | [N.D. - 0.24] |
| Country park | 9 | N.D. | N.D. | [N.D. - N.D.] | N.D. | N.D. | [N.D. - N.D.] | N.D | N.D. | [N.D. - N.D.] | N.D. | N.D. | [N.D. - N.D.] | N.D. | N.D. | [N.D. - N.D.] |
| Rural area | 19 | 0.04 | N.D. | [N.D. - 0.12] | 0.07 | 0.1 | [N.D. - 0.3] | 0.02 | N.D. | [N.D. - 0.11] | 0.13 | 0.1 | [N.D. - 0.48] | 0.02 | 0.02 | [N.D. - 0.09] |
| Restored landfill | 11 | 0.04 | N.D. | [N.D. - 0.12] | 0.09 | 0.1 | [N.D. - 0.24] | 0.01 | N.D. | [N.D. - 0.06] | 0.14 | 0.1 | [N.D. - 0.42] | 0.02 | 0.01 | [N.D. - 0.07] |
| Agricultural farmland | 9 | 0.05 | N.D. | [N.D. - 0.2] | 0.06 | N.D. | [N.D. - 0.25] | 0.02 | N.D. | [N.D. - 0.06] | 0.13 | N.D. | [N.D. - 0.47] | 0.02 | N.D. | [N.D. - 0.07] |
| Orchard farm | 5 | 0.05 | N.D. | [N.D. - 0.12] | 0.04 | N.D. | [N.D. - 0.12] | N.D | N.D. | [N.D. - N.D.] | 0.1 | 0.1 | [N.D. - 0.25] | 0.01 | 0.01 | [N.D. - 0.03] |
| Crematorium | 10 | 0.03 | N.D. | [N.D. - 0.11] | 0.05 | 0.0 | [N.D. - 0.11] | N.D | N.D. | [N.D. - N.D.] | 0.09 | 0.1 | [N.D. - 0.22] | 0.01 | 0.01 | [N.D. - 0.03] |
| Industrial area | 18 | 0.04 | N.D. | [N.D. - 0.22] | 0.07 | N.D. | [N.D. - 0.43] | 0.01 | N.D. | [N.D. - 0.1] | 0.13 | N.D. | [N.D. - 0.66] | 0.02 | N.D. | [N.D. - 0.08] |
| Nearby Highway | 4 | 0.16 | 0.1 | [N.D. - 0.44] | 0.51 | 0.3 | [N.D. - 1.53] | 0.16 | 0.1 | [N.D. - 0.51] | 0.83 | 0.4 | [N.D. - 2.48] | 0.15 | 0.06 | [N.D. - 0.45] |

Table 1 (con't)
| Classified soil categories | Sample no. | PCB-28 (NDL) |  |  | PCB-52 (NDL) |  |  | PCB-101 (NDL) |  |  | †PCB-118 (NDL) |  |  | PCB-138 (NDL) |  |  |
| --- | --- | --- | --- | --- | --- | --- | --- | --- | --- | --- | --- | --- | --- | --- | --- | --- |
|  |  | M* | Medi* | Range | M* | Medi* | Range | M* | Medi* | Range | M* | Medi* | Range | M* | Medi* | Range |
| Concentration in soils (µg/kg) |  |  |  |  |  |  |  |  |  |  |  |  |  |  |  |  |
| Urban park | 39 | 0.33 | 0.2 | [0.09 - 5.19] | 0.5 | 0.5 | [N.D. - 0.98] | 0.16 | 0.1 | [N.D. - 2.45] | 0.18 | 0.1 | [N.D. - 1.05] | 1.61 | 0.3 | [N.D. - 13.89] |
| Greening area | 14 | 0.36 | 0.2 | [N.D. - 2.49] | 0.66 | 0.4 | [0.09 - 2.8] | 0.14 | 0.1 | [N.D. - 0.91] | 0.2 | 0.1 | [N.D. - 1.24] | 0.39 | 0.3 | [N.D. - 1.52] |
| Country park | 9 | 0.13 | 0.1 | [0.07 - 0.39] | 0.41 | 0.4 | [0.24 - 0.76] | N.D. | N.D. | [N.D. - N.D.] | N.D. | N.D. | [N.D. - N.D.] | 0.04 | N.D. | [N.D. - 0.12] |
| Rural area | 19 | 0.23 | 0.2 | [N.D. - 1.42] | 0.48 | 0.5 | [N.D. - 1.2] | 0.06 | 0.1 | [N.D. - 0.24] | 0.07 | 0.1 | [N.D. - 0.3] | 0.32 | 0.3 | [N.D. - 1.03] |
| Restored landfill | 11 | 0.19 | 0.2 | [0.1 - 0.3] | 0.57 | 0.6 | [0.26 - 0.89] | 0.07 | N.D. | [N.D. - 0.29] | 0.09 | 0.1 | [N.D. - 0.24] | 0.29 | 0.1 | [N.D. - 1.11] |
| Agricultural farmland | 9 | 0.14 | 0.1 | [0.07 - 0.32] | 0.44 | 0.4 | [0.28 - 0.69] | 0.04 | N.D. | [N.D. - 0.15] | 0.06 | N.D. | [N.D. - 0.25] | 0.18 | 0.1 | [N.D. - 0.62] |
| Orchard farm | 5 | 0.13 | 0.2 | [0.07 - 0.17] | 0.37 | 0.4 | [0.32 - 0.41] | 0.04 | N.D. | [N.D. - 0.08] | 0.04 | N.D. | [N.D. - 0.12] | 0.15 | 0.1 | [N.D. - 0.34] |
| Crematorium | 10 | 0.16 | 0.1 | [0.06 - 0.51] | 0.49 | 0.4 | [0.31 - 0.73] | 0.05 | N.D. | [N.D. - 0.19] | 0.05 | 0.0 | [N.D. - 0.11] | 0.28 | 0.2 | [0.13 - 0.88] |
| Industrial area | 18 | 0.23 | 0.1 | [N.D. - 1.86] | 0.38 | 0.4 | [N.D. - 1.03] | 0.05 | N.D. | [N.D. - 0.36] | 0.07 | N.D. | [N.D. - 0.43] | 0.16 | 0.1 | [N.D. - 0.64] |
| Nearby Highway | 4 | 0.5 | 0.2 | [0.09 - 1.58] | 2.34 | 0.7 | [0.42 - 7.47] | 0.36 | 0.2 | [N.D. - 1.02] | 0.51 | 0.3 | [N.D. - 1.53] | 1.03 | 1.0 | [0.11 - 1.91] |
| Classified soil categories | Sample no. | PCB-153 (NDL) |  |  | PCB-180 (NDL) |  |  | Total 6 NDL PCBs |  |  | Total 7 NDL PCBs |  |  | Total 66 PCB congeners |  |  |
|  |  | M* | Medi* | Range | M* | Medi* | Range | M* | Medi* | Range | M* | Medi* | Range | M* | Medi* | Range |
| Concentration in soils (µg/kg) |  |  |  |  |  |  |  |  |  |  |  |  |  |  |  |  |
| Urban park | 39 | 0.94 | N.D. | [8.28 - 0.22] | 1.21 | 0.2 | [N.D. - 9.52] | 4.76 | 1.4 | [0.22 - 33.77] | 4.94 | 1.5 | [0.23 - 34.58] | 6.44 | 5.0 | [3.21 - 25.42] |
| Greening area | 14 | 0.27 | N.D. | [1.04 - 0.22] | 0.22 | 0.2 | [N.D. - 0.49] | 2.04 | 1.3 | [0.15 - 9.15] | 2.24 | 1.5 | [0.16 - 10.39] | 5.87 | 5.2 | [0.93 - 14.73] |
| Country park | 9 | 0.09 | N.D. | [0.35 - N.D.] | 0.04 | N.D. | [N.D. - 0.11] | 0.73 | 0.7 | [0.38 - 1.26] | 0.74 | 0.7 | [0.39 - 1.27] | 4.01 | 3.7 | [2.79 - 6.31] |
| Rural area | 19 | 0.24 | N.D. | [1.04 - 0.2] | 0.31 | 0.2 | [N.D. - 1.41] | 1.65 | 1.4 | [0.39 - 4.35] | 1.72 | 1.5 | [0.43 - 4.65] | 4.67 | 3.9 | [2.13 - 13.52] |
| Restored landfill | 11 | 0.27 | N.D. | [0.83 - 0.25] | 0.21 | 0.1 | [N.D. - 0.84] | 1.61 | 1.3 | [0.73 - 4.06] | 1.7 | 1.4 | [0.74 - 4.29] | 4.49 | 4.4 | [2.84 - 5.92] |
| Agricultural farmland | 9 | 0.24 | N.D. | [0.81 - 0.08] | 0.1 | N.D. | [N.D. - 0.51] | 1.13 | 0.9 | [0.46 - 2.71] | 1.19 | 0.9 | [0.47 - 2.97] | 3.84 | 3.7 | [2.39 - 5.12] |
| Orchard farm | 5 | 0.09 | N.D. | [0.24 - N.D.] | 0.09 | 0.1 | [N.D. - 0.23] | 0.86 | 0.6 | [0.53 - 1.38] | 0.91 | 0.6 | [0.54 - 1.5] | 5.09 | 4.0 | [3.79 - 9.84] |
| Crematorium | 10 | 0.25 | 0.1 | [0.83 - 0.17] | 0.22 | 0.2 | [0.06 - 0.74] | 1.45 | 1.4 | [0.87 - 3.01] | 1.5 | 1.5 | [0.88 - 3.12] | 3.48 | 3.2 | [2.18 - 5.64] |
| Industrial area | 18 | 0.1 | N.D. | [0.39 - N.D.] | 0.09 | 0.0 | [N.D. - 0.32] | 1.01 | 0.8 | [N.D. - 4.16] | 1.07 | 0.8 | [N.D. - 4.29] | 33.13 | 7.1 | [0.22 - 154.11] |
| Nearby Highway | 4 | 0.63 | 0.1 | [1.11 - 0.67] | 0.44 | 0.4 | [0.15 - 0.8] | 5.3 | 3.1 | [1.17 - 13.89] | 5.81 | 3.3 | [1.18 - 15.43] | 24.78 | 8.7 | [7.34 - 74.49] |

Values in square brackets represent range of concentration. \*M=Mean, Medi=Median. †: Not included for total 6 NDL PCBs. ‡: Calculated from sum of PCB 105, 118 and 157.

To assess the spatial distribution of individual DDTs in Hong Kong, kriging maps (Figure 1) were constructed by interpolating the data from various locations (ordinary kriging). A single hotspot for DDT, DDD and DDE contamination was observed in the southern tip of Tsing Yi island. Another hotspot for DDT compounds was found in the roadside located in Lung Kwu Tan, where no DDD and DDE were detected. The half life of DDTs ranges from a few months to 30 years, or even up to centuries (Aigner *et al.*, 1998; Dimond and Owen, 1996). The traffic in the aforementioned locations was mainly contributed by trucks powered by diesel engine, which may release high levels of toxic pollutants such as PAHs and lead, creating an adverse condition which is unfavorable to biodegradation, and thus slow down the biodegradation of DDTs in the vicinity.

**Figure 1.**
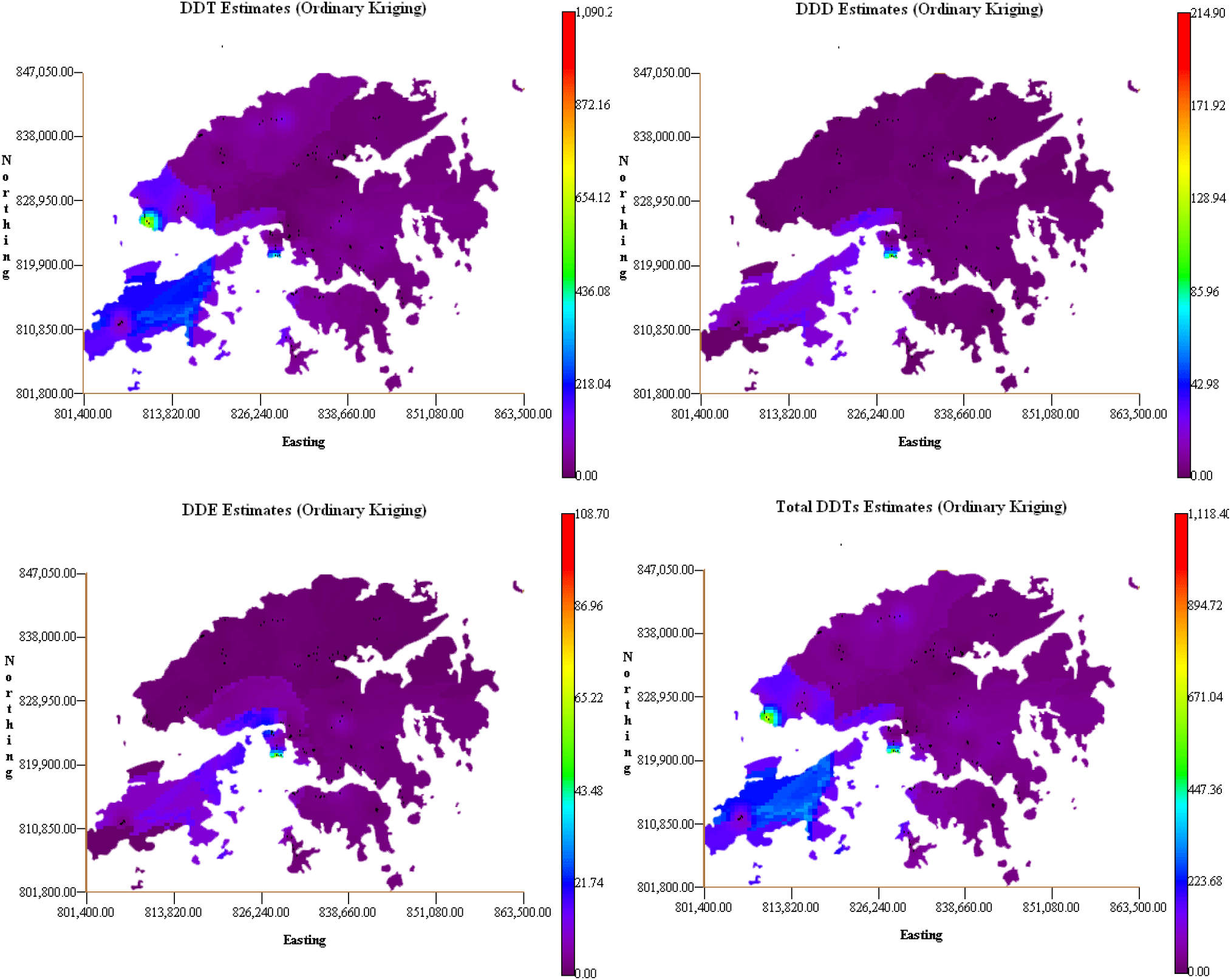
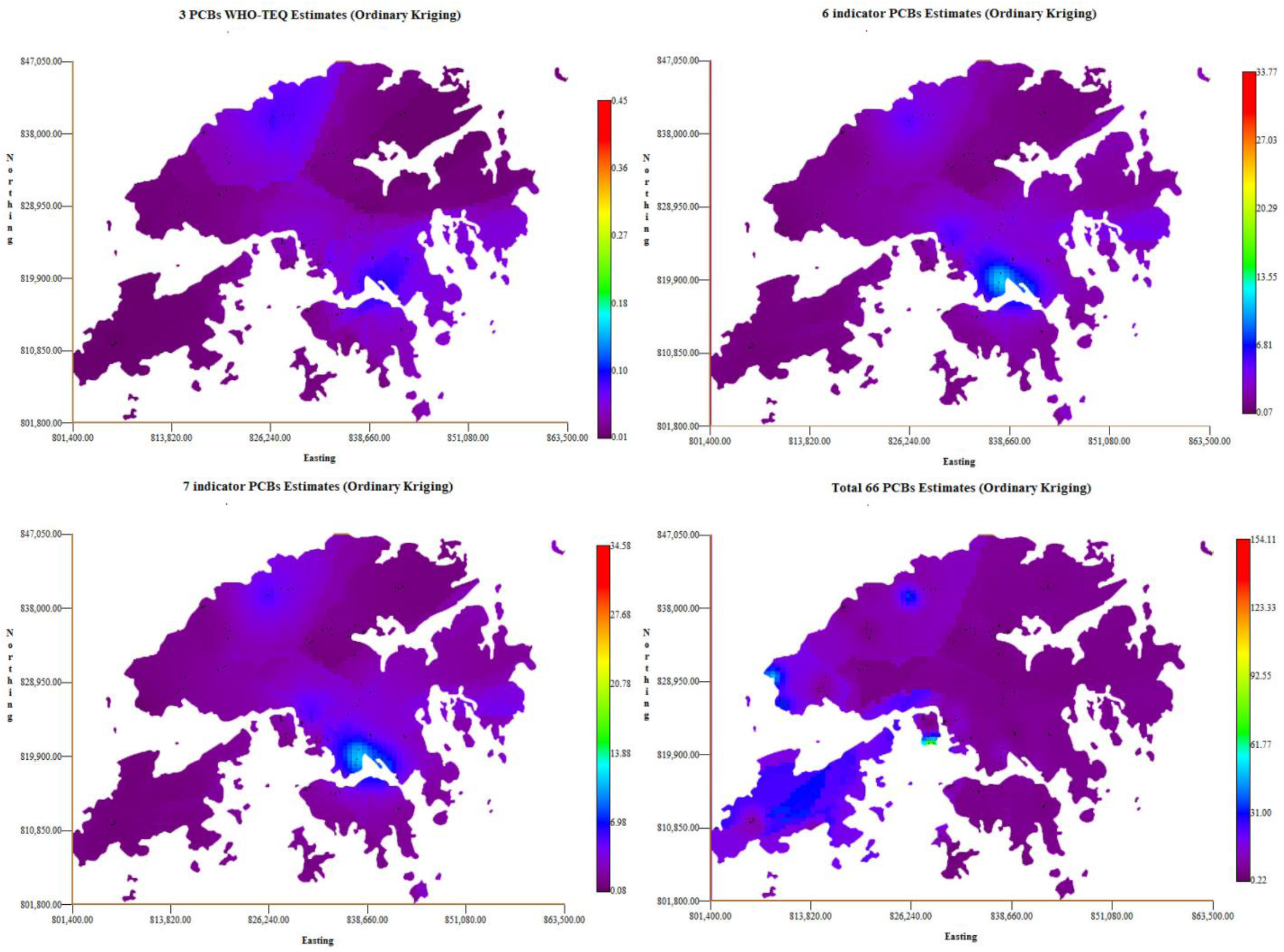
Kriged maps DDT (DDT, DDD, DDE and total DDTs) and PCB (3-PCBs WHO-TEQ congeners, 6 indicator PCBs, 7 indicator PCBs and sum of 66 PCB congeners) levels (µg/kg) in surface soils of Hong Kong. The 3-PCBs WHO-TEQ is in ng kg^−1^ scale.

In Hong Kong, relatively high levels of DDTs (2.87 mg kg^−1^ fat) and β-HCH (0.95 mg kg^−1^ fat) were found in human milk (Wong *et al.*, 2002), whereas the mean concentration of DDTs in whale and dolphin found in local water was 46 mg kg^−1^ wet wt. (Minh *et al.*, 1999). The present results indicated that the mean edaphic DDT concentrations was 72 µg kg^−1^, while levels of DDTs in river sediments and tilapia of the inland water systems in Hong Kong were 2.8 to 8.6 and 28 to 40 µg kg^−1^ dry wt. respectively (Zhou *et al.*, 1999), and average DDT levels in surface sediment of Mai Po marsh was 8.15 µg kg^−1^ dry wt (Zheng *et al.*, 2000). It can be seen that there are up to 1000 folds of difference in DDT concentrations between the backgrounds and living organisms, and this implied that DDTs are being accumulated into organisms and finally reaching human beings through various pathways including ingestion of contaminated food and soil particles, and inhalation of dust. Being a large agricultural country, China has produced and used a large quantity of DDTs not until early 1980s when DDT was banned for agriculture application (Wong *et al.*, 2005). The level of DDTs in China is highly varied, from maximum values below 100 µg kg^−1^ in crop soils of PRD (5 to 80 µg kg^−1^) (Fu *et al.*, 2003), to below 1000 µg kg^−1^ in Tianjin (0.35 to 963 µg kg^−1^) (Gong *et al.*, 2004) and above 1000 µg kg^−1^ in outskirts of Beijing (0.77 to 2178 µg kg^−1^) (Zhu *et al.*, 2005). Large difference between the minimum and maximum values is also observed in the current study (Table 1).

The concentrations of total PCBs, dioxin like (DL) and non-dioxin like (NDL) PCBs in soil samples are shown in Table 1. Concentrations of total PCBs could be grouped into 3 categories: The most contaminated which included industrial and areas near highway (median concentration around 8 µg kg^−1^); followed by urban park and greening area (5 µg kg^−1^) and the rest of the land uses which contained the lowest concentration (3 to 4 µg kg^−1^). The PCB toxicity is expressed as toxic equivalent quantity (TEQ) and is calculated according to toxic equivalency factor (TEF) stated by World Health Organization (WHO). It represents the total toxicity of a mixture of related substances that equal to their combined toxic effects. Median PCB WHO-TEQs (calculated from 3 DL PCBs) in all the land use categories were below 0.06 ng kg^−1^. Zhao *et al.* (2006) reported that the PCB WHO-TEQs (9 DL PCBs) of soils collected from a polluted and abandoned farmland in southern China was 57 ng kg^−1^, whereas the PCB WHO-TEQs equaled to 5.13 ng kg^−1^, which is much higher than the corresponding data of the present study. Among the 3 DL-PCBs, PCB-157 was contributing greater than 40% in all land use categories, followed by PCB-118 and PCB-105. For the sum of 6 and 7 NDL-PCBs, PCB-52 was the most dominant congener (usually greater than 35% of total NDL-PCBs), while PCB-101 was the lowest among the indicator PCBs. It was reported that ∑ 13 PCBs ranged from 0.051 to 22 µg kg^−1^ in soils of United Kingdom and Norway (Meijer *et al.*, 2002), while concentration of ∑7 PCBs ranged from 0.09 to 150 µg kg^−1^ (Seine River basin) in France (Motelay-Massei *et al.*, 2004). In addition, ∑6 PCBs in Moscow soils were found between 2 to 34 µg kg^−1^ (Wilcke *et al.*, 2006). PCBs in soils from Shenyang, China ranged from 6.4 to 15.2 µg kg^−1^, but the polluted areas in southeast coast of China where PCBs related activities such as dismantling of PCB-containing transformers reached 788 µg kg^−1^ (Xing *et al.*, 2005). The PCB concentrations in Hong Kong soils were within the lower range of the levels reported, because of a lack of PCB related industry.

### 3.2 Sources

Correlation matrixes for DDTs in different land use categories are presented in Table 2. In most cases DDD were significantly correlated to DDE (p<0.05). Since DDT could be degraded to DDE and then DDD in the environment, or directly to DDD (Agency for Toxic Substances and Disease Registry, 2002), the positive correlation of DDD and DDE suggested that DDE to DDD was not an important pathway for DDT degradation. The results also showed that DDT had a significant correlation with its metabolites in agricultural farmlands, industrial areas and restored landfills, thus a slow but continuous input of DDT to these areas is suspected, as increase in metabolites should be accompany with a decrease in parent compound.

**Table 2.**
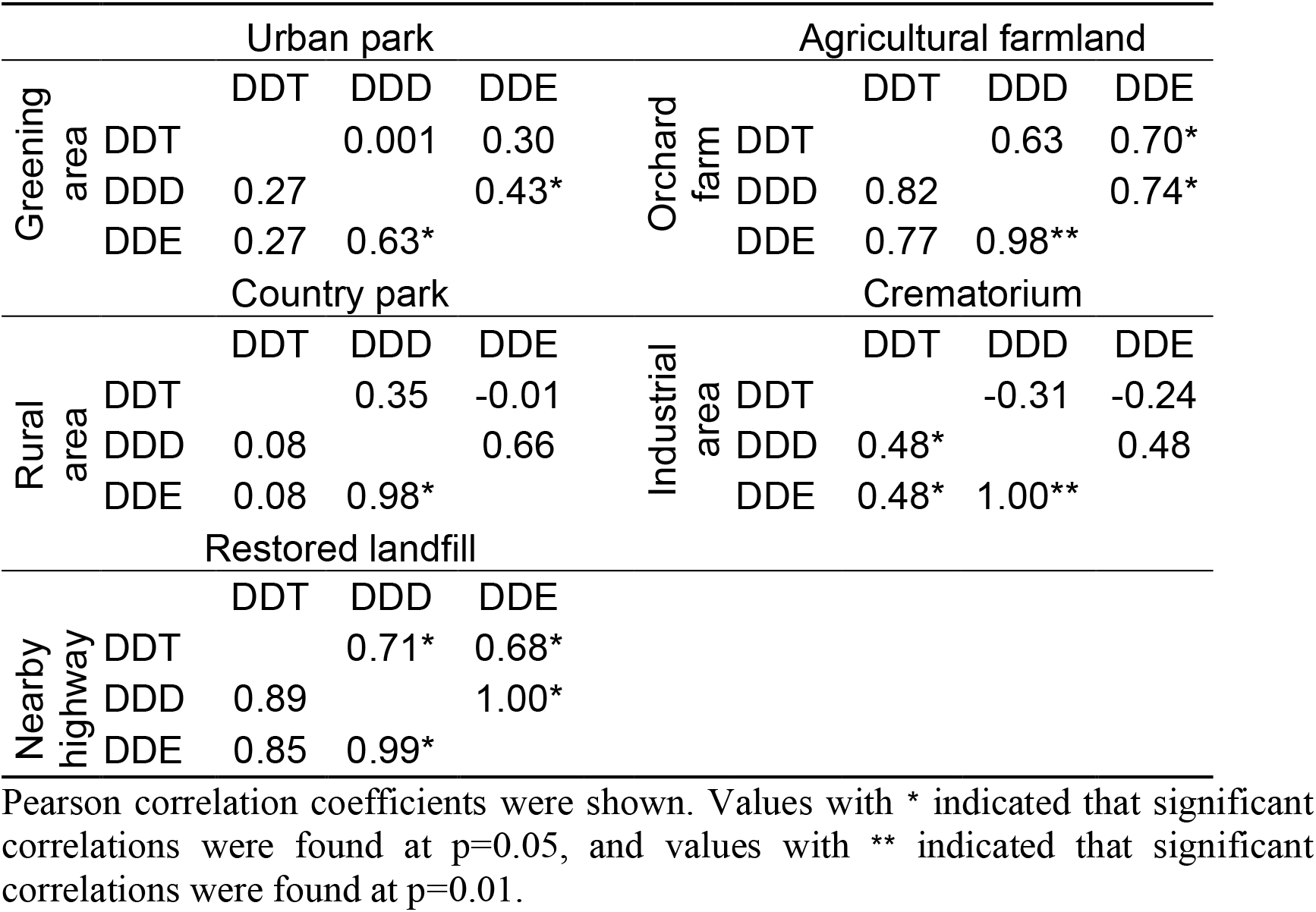
Correlation matrix of DDT concentrations for each individual land use in Hong Kong.

|  |  | Urban park |  |  |  |  | Agricultural farmland |  |  |
| --- | --- | --- | --- | --- | --- | --- | --- | --- | --- |
| Greening area |  | DDT | DDD | DDE | Orchard farm |  | DDT | DDD | DDE |
|  | DDT |  | 0.001 | 0.30 |  | DDT |  | 0.63 | 0.70* |
|  | DDD | 0.27 |  | 0.43* |  | DDD | 0.82 |  | 0.74* |
|  | DDE | 0.27 | 0.63* |  |  | DDE | 0.77 | 0.98** |  |
|  |  | Country park |  |  |  |  | Crematorium |  |  |
| Rural area |  | DDT | DDD | DDE | Industrial area |  | DDT | DDD | DDE |
|  | DDT |  | 0.35 | -0.01 |  | DDT |  | -0.31 | -0.24 |
|  | DDD | 0.08 |  | 0.66 |  | DDD | 0.48* |  | 0.48 |
|  | DDE | 0.08 | 0.98* |  |  | DDE | 0.48* | 1.00** |  |
|  |  | Restored landfill |  |  |  |  |  |  |  |
| Nearby highway |  | DDT | DDD | DDE |  |  |  |  |  |
|  | DDT |  | 0.71* | 0.68* |  |  |  |  |  |
|  | DDD | 0.89 |  | 1.00* |  |  |  |  |  |
|  | DDE | 0.85 | 0.99* |  |  |  |  |  |  |

Pearson correlation coefficients were shown. Values with \* indicated that significant correlations were found at $p=0.05$ , and values with \*\* indicated that significant correlations were found at $p=0.01$ .

Various DDT ratios could be used to assess any recent inputs of DDT in our environment (Kong *et al.*, 2005). These include DDD/DDT, DDE/DDT and DDT/(DDE+DDD) ratios. Table 3 presents the various calculated ratios for DDTs in each soil land use category. Metabolite(s) (DDD, DDE, DDE+DDD) to parent compound (DDT) ratios were less than 0.15 on average. In contrast, DDT/(DDE+DDD) ratios were much higher, particular in the vicinity of highways. Two hot spots (in Tsing Yi island, 796 µg kg^−1^ and Lung Kwu Tan, 1090 µg kg^−1^) identified in the kriged map (Figure 1) were both located adjacent to industrial activities, e.g. power station/oil depot/chemical plants/machinery factories in Tsing Yi island, and steel mill/power station/cement plant in Lung Kwu Tan. Out of the 138 soil samples collected, 18% of them did not show detectable level of neither DDT nor its metabolites. In addition, DDT without metabolites was detected in 25% of the 138 samples. These results suggested 2 possibilities: 1) there was still a small-scale, localized non-mobile DDT input or 2) the highly heterogeneous soil micro-environment could either promote degradation of DDT or it could inhibit the degradation of organic compounds, depending on the sampling locations. Photo-oxidation and volatilization of DDT are possible mechanisms for the loss of DDT (Hussain *et al.*, 1994), but the principal pathway for the loss of DDT in soils is mainly through microbial actions (Mohn and Tiedje, 1992).

**Table 3.** Various ratios for DDT and its metabolites calculated for each land use in Hong Kong soil.

| Land uses | DDD / DDT | DDE / DDT | DDT / (DDD+DDE) |
| --- | --- | --- | --- |
| Urban park | 0.17 ± 0.30 (a) | 0.08 ± 0.08 (a) | 6.33 ± 5.96 (a) |
| Greening area | 0.15 ± 0.10 (a) | 0.09 ± 0.08 (a) | 4.35 ± 6.54 (a) |
| Country park | 0.07 ± 0.10 (a) | 0.02 ± 0.05 (a) | 8.38 ± 7.14 (a) |
| Rural area | 0.06 ± 0.08 (a) | 0.03 ± 0.04 (a) | 9.45 ± 7.67 (a) |
| Restored landfill | 0.38 ± 0.19 (a) | 0.20 ± 0.11 (ab) | 0.80 ± 1.21 (a) |
| Agricultural farmland | 0.12 ± 0.11 (a) | 0.12 ± 0.07 (a) | 6.92 ± 5.07 (a) |
| Orchard farm | 0.16 ± 0.11 (a) | 0.36 ± 0.66 (b) | 3.56 ± 2.76 (a) |
| Crematorium | 0.10 ± 0.10 (a) | 0.05 ± 0.05 (a) | 4.79 ± 4.00 (a) |
| Industrial area | 0.20 ± 0.19 (a) | 0.08 ± 0.09 (a) | 4.84 ± 5.92 (a) |
| Nearby highway | 0.01 ± 0.01 (a) | 0.01 ± 0.01 (a) | 58.73 ± 42.14 (b) |

Values in parenthesis represent grouping by one way ANOVA analysis (Tukey HSD, $p < 0.05$ ).

The current official use of DDT is for the control of disease vectors in indoor house spraying as specified by the World Health Organization (WHO) (World Wildlife Fund, 2005). Apart from DDT, *p,p’*-DDD was also used as an insecticide but it is not as common as DDT (Agency for Toxic Substances and Disease Registry, 2002).

In Hong Kong, DDT is not a registered pesticide and has been banned for use since the beginning of 1988. Currently, it can be traded only under permit in Hong Kong (UNEP Chemicals, 2002). Between 1979 to 1982, about 5,023 to 5,996 kg of DDT pesticide was imported annually (Morton, 1990) and the existence of residual DDT and its metabolites in soils are therefore expected.

OCPs are routinely found in the atmosphere (Cortes *et al.*, 1998). A recent study observed that OCPs like α-HCH, hexachlorobenzene, DDT, DDE, heptachlor, and endosulfan I were detected in local atmospheric compartment at relatively low concentrations 0.02–0.23 ng m-3 (Louie and Sin, 2003). It was proved that the northeast monsoon wind can bring air pollutants from China to Hong Kong (Lee and Hills, 2003), and it is reasonable to deduce that pollutants such as DDTs bound in fine particles could also be a potential source in Hong Kong soils via atmospheric deposition of dust particles from the PRD.

Dicofol is an organochlorine acaricide (a chemical that kills mites) which is highly toxic to aquatic life and can cause egg-shell thinning in some bird species. As Dicofol is manufactured from technical DDT in China, it also contributes to the sources of fresh DDT in the surrounding (Qiu *et al.*, 2005). Though DDT is also released to the environment by pyrogenic degradation of animal fat in food during cooking in PRD, but its contribution to the edaphic environment is expected to be comparatively low (Cheng *et al.*, 2000). In addition, some OCPs (such as DDT and HCH) may still be illegally used for agricultural purpose in the PRD (Wong and Poon, 2003).

PCB homologues profile for corresponding land use categories is plotted in Figure 2. PCB patterns were not characterized in terms of lower or higher chlorinated biphenyl, but by individual homologues. Di-PCBs was the most dominant homologue, together with tetra- and octa-PCBs, they accounted for more than 70% of all the investigated PCBs. Aroclor is a commercial mixture of PCBs that has been used intensively. Homologue composition of Arochlor such as Arochlor 1016, 1242, 1254 and 1260 showed domination of tri-to hexa-PCBs (U.S. Environmental Protection Agency, 2005). However, the present results revealed that the major homologue was di-PCBs and thus rule out the possibility of recent Arochlor contamination. Interestingly, the contribution of octa-PCB accounted for more than 40% of total PCBs found in industrial areas. The greater distribution of heavier homologues is possibly due to the preferential atmospheric deposition in the vicinity of the source (Meijer *et al.*, 2002). It has been reported that combustion process such as municipal solid waste incineration, automobile exhaust are potential sources of PCBs (Granier and Chevreuil, 1991; Mai *et al.*, 2005) and the industrial activities, with mobile trucks could be the major contributor to PCBs in industrial areas.

**Figure 2.**
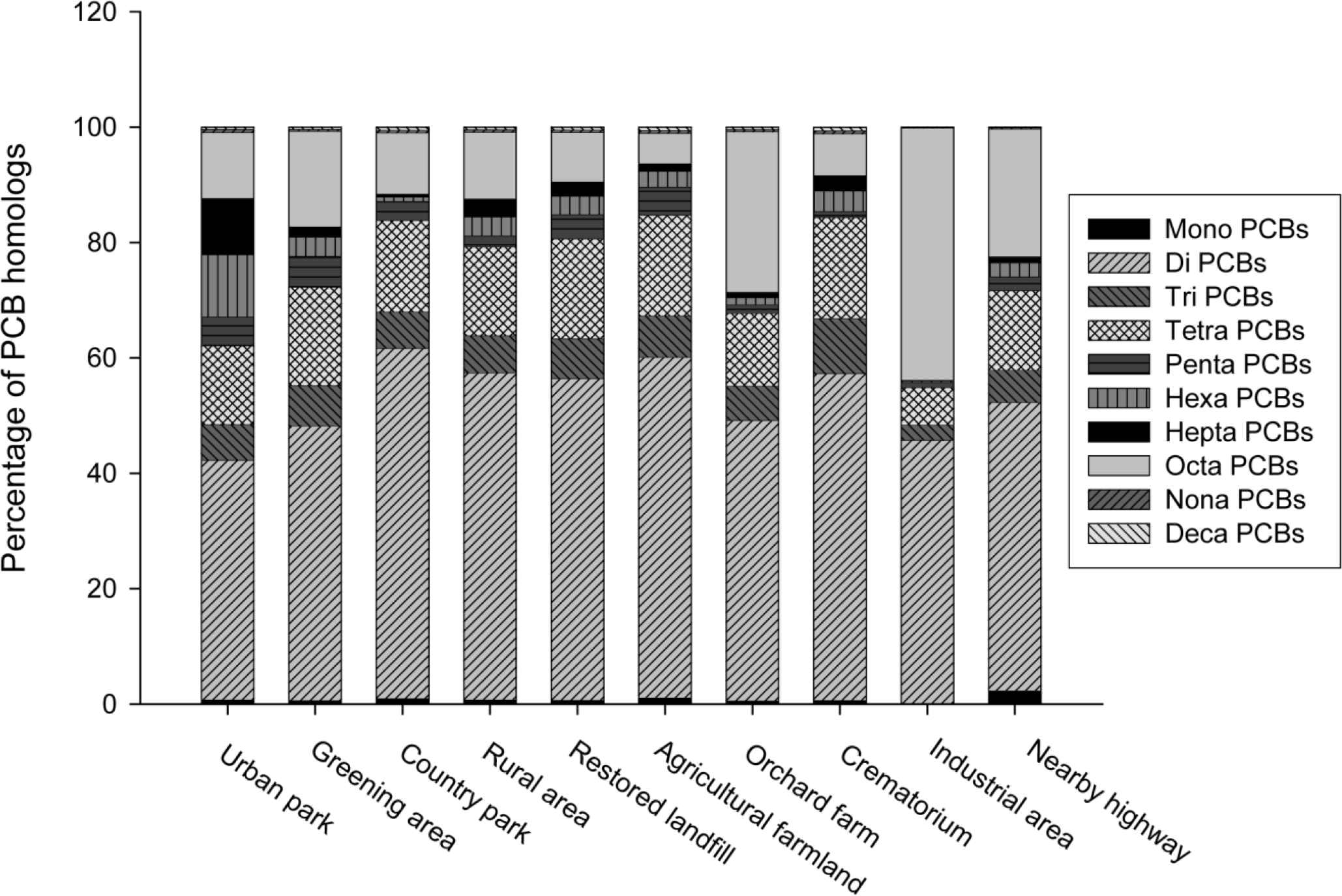
Homologue profiles of PCBs in corresponding land uses in Hong Kong. Total PCBs is sum of 66 selected congeners.

The general domination of di-PCB contaminations could be originated from the short-range atmospheric transport from industrial sites (Motelay-Massei *et al.*, 2004), and medium-range regional atmospheric deposition is also suspected (Wilcke *et al.*, 2006). Volatization and re-deposition during dredging of contaminated sediments is also a possible source of PCBs in soils (Vorhees *et al.*, 1999). However, a study indicated that total PCBs (∑112 congeners) in the sediments of PRD was in the range of 26 to 32 µg kg^−1^, with tri-to hexa-PCBs accounted for more than 80% of total PCBs (Mai *et al.*, 2005), thus this volatization and re-deposition pathway from sediment is unlikely to play a crucial role in PCB contaminations in soils.

### 3.3 Cleanup guidelines

The guideline values for DDTs and PCBs from various countries are listed in Table 4. In general, concentrations of DDT obtained in the present study were below the soil quality limits imposed by Netherlands (Ministry of Housing Spatial Planning and Environment, 2000), Denmark (Danish Environmental Protection Agency, 2002), Canada (Environment Canada, 2003) and China (State Environmental Protection Administration of China, 1995) except samples from southern Tsing Yi island (796 µg kg^−1^) and Lung Kwu Tan (1090 µg kg^−1^). For total DDTs (DDT + DDE + DDD), only one sample from Lung Kwu Tan (1090 µg kg^−1^) was slightly above of the guideline values from Denmark (1000 µg kg^−1^) while the rest of 137 samples were below the safety values from Netherlands, Denmark and China. In contrast to the sporadic occurrence of hazardous samples concerning DDTs, none of the samples was found to contain PCBs at alarming concentration when benchmarking with the soil quality guidelines from the aforementioned countries. In a word, the potential risks to human and ecosystem imposed by the investigated OC chemicals are minimum.

**Table 4.** Soil quality guidelines and their criteria for DDTs and PCBs in various countries.

| Country of implementation | Quality guidelines | Concentration in soils (µg/kg) |  |  |  |  |
| --- | --- | --- | --- | --- | --- | --- |
|  |  | DDT | DDD | DDE | Total DDTs | PCBs |
| Netherlands | Dutch target value | N.A. | N.A. | N.A. | 10* | 20‡ |
|  | Dutch intervention value | N.A. | N.A. | N.A. | 4000* | 1000‡ |
| Denmark | Danish soil quality criteria | N.A. | N.A. | N.A. | 1000# | N.A. |
| Canada | Canadian environmental quality guidelines | 700† | N.A. | N.A. | N.A. | (500)<br>1300† |
| China | Environmental quality standard | N.A. | N.A. | N.A. | N.A. | N.A. |

\* Sum of DDT, DDD and DDE.

## 4. Conclusions

In the present study of DDT and PCB concentration in Hong Kong soils, sporadic contamination of DDTs and it’s metabolites were found at a range of 0.22 to 154 µg kg^−1^ and the concentration was found to be highest in industrial areas. There were widespread but low levels of PCB contamination (0.22 to 154 µg kg^−1^), while the highest level was observed in industrial areas. PCB-157 was the most active DL-PCBs among 3 (PCB −105, 118, and 157) that have been investigated and contributed more than 40% of WHO-TEQ in all land use categories, while PCB-152 was the most dominant congener among the 6 or 7 indicator PCBs. The current levels of DDT and PCB in Hong Kong were within the guideline values of the soil quality guidelines adopted in countries such as Netherlands and Denmark. However, 3 out of 138 samples exceeded DDT(s) concentrations but their potential hazards were minimal as the sites were located in remote areas. It is advised that a thorough site investigation on DDTs should be made prior to any residential development in these locations.

## Acknowledgments

The authors are grateful to Mr. Y.Y. Chin from Leisure and Cultural Services Department (LCSD, HKSAR) for providing technical assistance. Financial supports from the Strategic Research Fund from the Science Faculty, HKBU and the Area of Excellence (AoE) Scheme under the University Grants Committee of the Hong Kong Special Administrative Region (CITYU/AoE/03-04/02) are also gratefully acknowledged.

